# fNIRS reveals that live social interactions and visual realism influence neural responses

**DOI:** 10.64898/2026.08.05.742785

**Authors:** Michaela Kent, Eva Deligiannis, Kevin M. Stubbs, Karsten Babin, Emma G. Duerden, Jody C. Culham

**Affiliations:** Neuroscience Program, University of Western Ontario, London, Ontario, Canada; Centre for Brain and Mind, University of Western Ontario, London, Ontario, Canada; Western Institute for Neuroscience, University of Western Ontario, London, Ontario, Canada; Department of Psychology, Faculty of Social Sciences, University of Western Ontario, London, Ontario, Canada; Faculty of Education, University of Western Ontario, London, Ontario, Canada

**Keywords:** fNIRS, social cognition, social interaction, faces, theory of mind

## Abstract

The human face is central to social interactions, supporting the ability to interpret others’ mental states using theory of mind (ToM). We examined whether functional near-infrared spectroscopy (fNIRS) would reveal brain-activation differences between live and pre-recorded social conversations in brain regions implicated in ToM. Furthermore, we examined whether activation depended on the visual realism of a social partner – viewed as a human or an animated avatar. By one view, social interactions may be dependent on how natural the social partner appears; by another view, social interactions may depend only upon the attribution of responses to a real human regardless of visual appearance. Neural activation for pre-recorded compared to live interactions was prolonged, consistent with extended cognitive effort. Activation patterns in the right temporoparietal junction differed between interacting with humans versus avatars, along with a stronger preference for looking at the eyes when interacting with a human (vs. avatar), underscoring the social relevance of real faces. Findings highlight the importance of both live interactions and facial realism in shaping social-cognitive processing, a finding with relevance for optimizing online social interactions.

## Introduction

In recent years, technological advances have transformed human social interactions. Since the COVID-19 pandemic, video conferencing has become ubiquitous, including both live meetings and pre-recorded presentations. In addition, the growing sophistication and use of artificial intelligence (AI) has driven rapid adoption of artificial agents, from cartoonish avatars (e.g., Apple’s Memojis) to highly realistic synthetic faces on the threshold of indistinguishability from real humans (“deep fakes”). Understanding people’s behaviour and brain responses across this realism continuum is therefore becoming increasingly important.

The growth of social neuroscience has enabled a better understanding of the neural substrates of human interactions, particularly through functional magnetic resonance imaging (fMRI) and functional near-infrared spectroscopy (fNIRS). FNIRS offers advantages for studying real-time interactions in more ecologically valid settings than fMRI permits (Schilbach *et al*. 2013, Redcay and Schilbach 2019) and is thus well-suited to examine how technologically mediated interactions engage, or attenuate, the social brain.

Natural social interactions rely on non-verbal cues such as eye gaze and speech timing to support the shared psychological state required for successful communication (Tomasello *et al*. 2005, Bailenson 2021). They also require the capacity to infer others’ mental states, referred to as mentalizing (Frith CD and Wolpert 2003, Frith U and Frith 2003) or theory of mind (ToM; Premack and Woodruff 1978). Understanding the neural underpinnings of these processes is especially relevant for conditions such as autism spectrum disorder (ASD), where face perception, perspective-taking, and ToM are often impaired (Hamilton, Brindley, and Frith 2009, Pedreño *et al*. 2017).

Neuroimaging has revealed a “social brain” network, including the medial prefrontal cortex (mPFC) and temporoparietal junction (TPJ), which is central to processing social information (Saxe and Kanwisher 2003, Adolphs 2009). Classic paradigms such as false-belief reasoning tasks (Saxe, Carey, and Kanwisher 2004, Dodell-Feder *et al*. 2011) and naturalistic stimuli like the cartoon film *Partly Cloudy* (Jacoby *et al*. 2016, Richardson *et al*. 2018) have been used to localise these regions. Importantly, however, the social brain’s response is shaped by the nature of the interaction itself. Gaze patterns differ when viewing a person one could interact with versus watching a video of the same person (Laidlaw *et al*. 2011, Risko *et al*. 2012, Freeth, Foulsham, and Kingstone 2013), and real-time, bidirectional exchanges elicit stronger and more synchronised activation across social brain regions than passive observation, reflecting the importance of mutual contingency and live responsiveness(Hirsch *et al*. 2018, Noah *et al*. 2020, Zhao *et al*. 2023). Relatedly, cross-brain neural coupling in the angular gyrus, a region closely associated with TPJ (Schurz *et al*. 2017), is specifically enhanced during real-time eye-to-eye contact compared with video-like gaze, even when the visual content is matched (Noah *et al*. 2020). Social brain responses also vary with perceived partner agency: reward-related circuits are more engaged when individuals believe they are interacting with a human, whereas an agent perceived as non-human or computer-controlled preferentially recruits attention and cognitive control networks (Pfeiffer *et al*. 2014).

A landmark step toward more ecologically valid social neuroimaging was taken by Redcay and colleagues (2010), who developed a method for simulating live face-to-face interactions during fMRI via a video feed. This approach demonstrated that genuinely contingent social exchanges, compared to non-interactive or recorded conditions, elicit increased activation in social brain regions, including the right TPJ and posterior superior temporal sulcus. Crucially, this work highlighted that contingent responding and joint attention – two hallmarks of real social interaction that are absent from conventional neuroimaging paradigms – are key drivers of social brain engagement.

Building directly on this work, subsequent studies asked whether similar effects emerge when contingency is manipulated at the level of belief rather than actual interaction. Rice et al. (2016; Rice and Redcay 2016) reported greater TPJ engagement when participants believed they were interacting in live compared to pre-recorded conditions, along with associations between autistic traits and activity in left angular gyrus. However, these effects were obtained under conditions in which “live” interactions were in fact pre-recorded and they lacked a video component, limiting conclusions about true real-time contingency and leaving open whether social brain responses reflect actual interaction dynamics or experimental belief alone.

The visual realism of the interaction partner also matters: interactions with an avatar controlled by another person support better learning than identical virtual agents without a human operator (Okita, Bailenson, and Schwartz 2007), and robot faces produce reduced ToM-region activation relative to human faces, particularly in right mPFC and TPJ (Gobbini *et al*. 2011). Yet, by one perspective, it may be beliefs and expectations about an agent, rather than its visual features per se, that most powerfully drive social brain engagement (Hortensius and Cross 2018).

Here we used fNIRS to investigate whether social brain regions respond more strongly to real-time interactions (live versus pre-recorded) and/or the visual realism of the partner (human versus avatar) during a video conversation. Figure 1 illustrates the experimental task, in which participants listened to a story told by a partner (confederate) and then answered a question requiring inference of the partner’s mental state (adapted from Rice, Moraczewski, and Redcay 2016). Eye tracking was recorded to assess potential gaze differences across conditions, and participants completed the Autism Spectrum Quotient (AQ) to examine whether autistic traits relate to neural responses during these interactions (Baron-Cohen *et al*. 2001).

**Figure 1.**
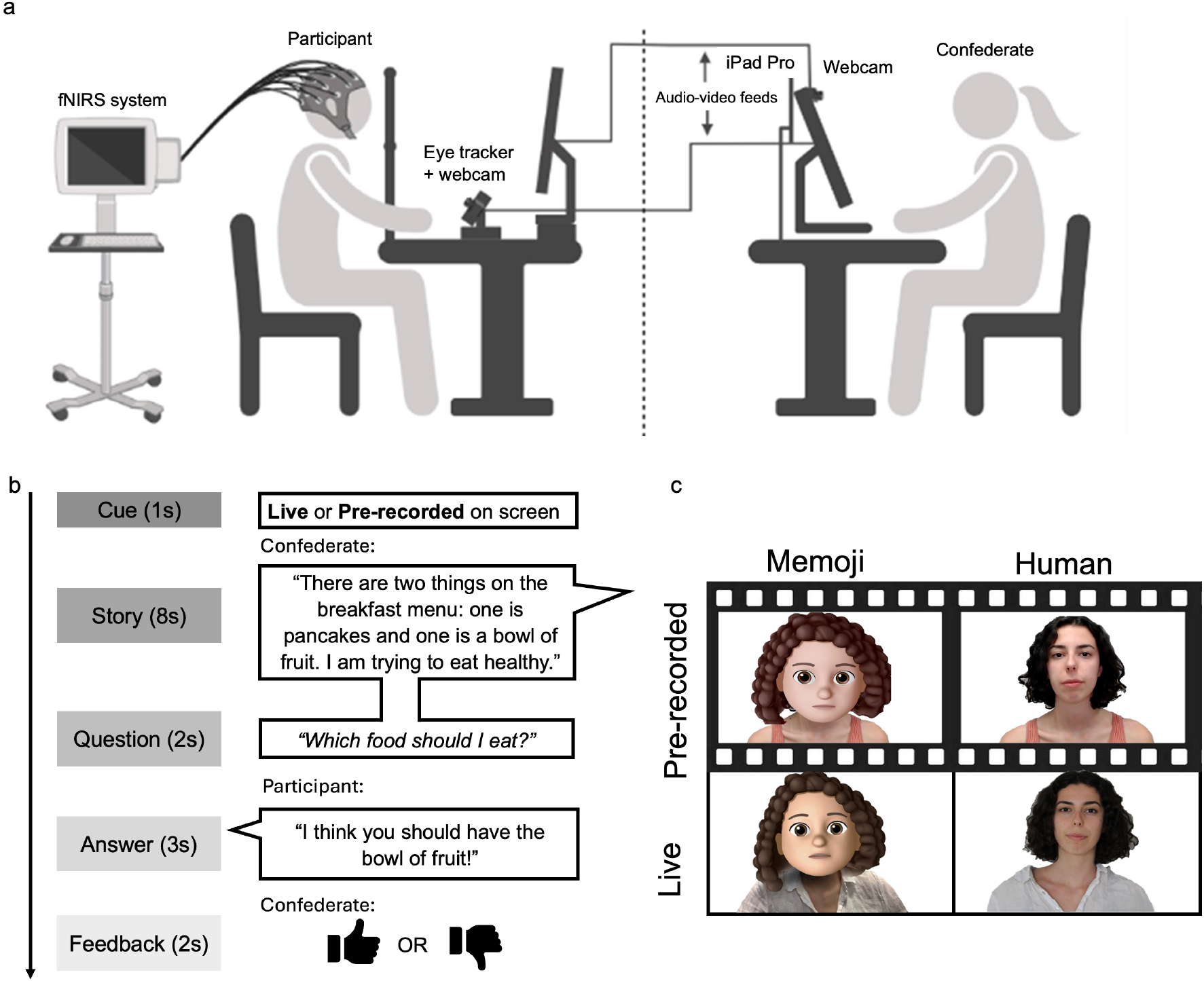
Experimental setup, stimuli and example trial timeline. (a) Experimental setup showing participant and confederate seated in front of their respective equipment required for two-way video and audio communication. The participant and confederate were in separate rooms for the duration of the task runs and could not see one another other than via the monitor. (b) Participants first received a cue as to whether the condition would be live or pre-recorded. The confederate explained a short story presenting the participant a situation with two options, indicating their preference, and then asking the participant which option they would prefer. Then the participant responded verbally with their choice and received positive or negative feedback from the confederate indicated by a thumbs-up or thumbs-down, respectively. (c) The four conditions showing manipulations of visual realism (human and avatar) and live and pre-recorded conditions. Note: The human face shown in (c) is that of author E.D., who has given express consent for these images to be published. The avatar was generated from the same individual. **Alt text:** Three-part schematic: (a) a participant in an fNIRS cap at a monitor with eye tracker and webcam, linked by two-way audio-video to a confederate in a separate room; (b) a trial timeline of cue, story, question, answer and feedback, annotated with an example exchange; (c) a two-by-two grid of Memoji and human faces, each shown live and pre-recorded.

Our primary aim was to investigate the neural responses to human versus avatar faces. Rather than comparing a real face to a highly realistic avatar, we used a cartoonish avatar (Apple “Memoji”) – clearly synthetic, yet controlled by a real human’s movements and expressions – to maximize any potential differences in neural responses. By one hypothesis (Hortensius and Cross 2018), neural and gaze responses may depend on perceived agency rather than visual realism. Alternatively, the absence of subtle emotional cues (e.g., facial microexpressions) in a cartoon avatar may attenuate social network recruitment relative to a natural face.

The secondary aim was to examine whether live versus pre-recorded interactions differentially engage ToM regions, as a conceptual replication of Rice et al. (2016) using fNIRS, with the key methodological improvement that our live interactions were truly live and we used visual stimuli throughout the interaction. We expected to replicate prior findings of greater TPJ activation for live interactions, and our 2×2 design allowed us to examine whether realism and liveness interact – potentially dissociating the most naturalistic condition (live human) from the three less naturalistic ones.

Finally, the third aim was to characterise the temporal dynamics of brain activation across different phases of the social interaction. Previous fMRI work (Rice, Moraczewski, and Redcay 2016) used a boxcar predictor, assuming uniform activation. However, mentalizing demands may increase over the story and extend into the question-and-answer phase. We therefore employed a deconvolution approach (finite impulse response functions; Glover 1999) to estimate activation time courses without assumptions about the hemodynamic response – an approach that has recently been implemented for fNIRS (Santosa *et al*. 2019, Machado *et al*. 2021) and has been shown to outperform conventional averaging by modelling trial-history confounds (Aarabi, Osharina, and Wallois 2017).

## Methods

### Participants

Data were analyzed from 25 participants (*n =* 13 males, 12 females; *M* age = 22 years). Data from an additional eight participants were collected but discarded because they did not pass initial fNIRS screening (*n* = 7) or due to insufficient fNIRS data (*n* = 1). The fNIRS screening protocol involved participants completing a short finger-tapping experiment to assess signal quality in channels covering the motor cortex. Data quality was monitored during acquisition using the NIRx recording software (NIRStar; NIRx Medical Technologies) by visually inspecting raw channel time series for noise, saturation, and signal dropout. Signal quality was quantified using the scalp coupling index (SCI), an automated NIRx metric indexing the consistency of cardiac pulsatility; channels with low SCI values, indicative of poor optode– scalp coupling, were flagged and adjusted when possible prior to data collection.

Participants were recruited using The University of Western Ontario’s Cognitive Neuroscience Research Registry (OurBrainsCAN.uwo.ca), the Western Psychology Research Participation Tool (SONA), and through advertisements. All individuals were native English speakers with normal or corrected-to-normal vision. Written informed consent was obtained prior to participation, and the study was approved by the university’s Non-Medical Research Ethics Board in accordance with the 1964 Declaration of Helsinki. Compensation was provided in the form of course credit or financial reimbursement (15 Canadian dollars per hour). Participants who were excluded after data quality screening were also compensated for their time.

### Functional Near-Infrared Spectroscopy

#### Stimuli

A video stream was manipulated to create four conditions: live human, live avatar, pre-recorded human, and pre-recorded avatar (Figure 1). Video was either presented to participants live or by replaying a pre-recorded video of a trial from a previous participant. Using previous live trials ensured that the confederate’s behaviour was consistent with a live interaction and reduced possible differences between live and pre-recorded conditions that may have confounded the results. The confederate’s face was either shown as their real face or as an avatar face using an Apple “Memoji”. All four conditions had the same voice and body; the only visual difference between the conditions was the face of the confederate and clothing. For the live conditions, the confederate was shown wearing the same shirt as they wore when meeting the participant in person before the experiment began; whereas, for the pre-recorded condition, the shirt was different.

For human conditions, the confederate video was obtained using Logitech webcams and microphone (Logitech C920). For avatar conditions, the video was obtained using an Apple iPad Pro (2020, iOS version 14.6) with built-in Memoji face filter. All trials were recorded and saved as video files for later analysis.

#### Pre-test procedure

The confederate met the participant in person, wearing the same clothing as in the live conditions to facilitate believability. After the confederate left the testing room, all communication was done by a live, two-way video call.

#### Test procedure

Participants engaged in a question-and-answer task, responding verbally to two-alternative forced-choice questions. The social interaction task was adapted from Rice and Redcay (2016) for use with fNIRS, with the addition of the live video call and avatar conditions. Importantly, the fNIRS version of this task allowed participants to respond verbally to more closely simulate a real social interaction. For all conditions, the trial structure remained the same (Figure 1b). The confederate gave a short vignette (story) before asking a question. The participant was asked to choose the “correct” option based on the information provided. For live conditions, the confederate gave a thumbs-up or thumbs-down response by video and for the pre-recorded conditions these responses were given with a static image. Each trial lasted 16 s followed by an inter-trial interval (ITI) of 6, 7, 8, or 9 s, with a higher preponderance of shorter ITIs. Each run began and ended with a 30-s baseline, during which participants were asked to remain still and fixate on a cross in the middle of the screen.

Participants completed four runs of sixteen trials (four trials per condition per run), with each run lasting approximately 8 minutes (calibration + 16 trials + ITIs + baselines). Placement of all conditions within the same run facilitates statistical comparisons among them because it avoids any differences in noise levels that can affect signal normalization. As such, considerable technical development was conducted to enable seamless alternation between human, avatar, live, and pre-recorded conditions within the same runs.

The order of conditions was counterbalanced using OptSeq2 (http://surfer.nmr.mgh.harvard.edu/optseq/) such that each condition was preceded an equal number of times by every other condition (including itself and the initial baseline period) to reduce potentially confounding effects of order history. The assignment of specific conversation topics to conditions was randomized for each participant. Trials were presented in a jittered event-related design to minimize the overlap of the hemodynamic response between multiple trials (Petersen and Dubis 2012) and to reduce contamination of systematic physiological noise such as Mayer waves and respiration (Yücel *et al*. 2016).

In total, the session lasted approximately 1.5 hours, including approximately 20 minutes for setup and instructions, 40 minutes for the main experiment, 20 minutes for localizers, and 10 minutes for questionnaires.

#### Partly Cloudy localizer

*Partly Cloudy* (Disney Pixar) is a 5.6-minute animated cartoon that contains moments recruiting ToM or mentalizing, as well as scenes that show the main character experiencing pain. Contrasting the mentalizing events with pain events (mental > pain) has been shown to identify the ToM network in fMRI (Jacoby *et al*. 2016, Richardson *et al*. 2018). The participant watched the movie passively and was asked to remain still for the duration.

#### False-belief task localizer

In the false-belief task, participants were asked to read false-belief stories (describing incorrect beliefs about the world), and false-photograph stories (describing outdated photographs, maps, or signs of the world). Each of the stories was followed by a statement about the story and participants were asked to indicate if the statement about the story was true or false using the keyboard. The stories were presented for 15 s, followed by the statements for 6 s, after which the participant was required to respond. This task took approximately 10 minutes to complete. The false belief events > false photo events contrast of interest has been shown to identify the ToM network in fMRI (Dodell-Feder *et al*. 2011).

#### Montage design

A custom montage was created following the international 10-5 EEG system for electrode placement (see supplementary Figure S1). Sensitivity modeling of the montage was carried out in AtlasViewer (Aasted *et al*. 2015) and the montage was created in NIRSite (NIRx Medical Technologies LLC, Berlin, Germany).

#### Data acquisition

FNIRS data were collected using a continuous wave NIRScout system (NIRx Medical Technologies LLC, Berlin, Germany). For the current study, 13 laser sources, 31 long-distance detectors (approximately 3 cm from source), and 4 short-distance detectors (0.8 cm from source) were used, resulting in 64 channels. Laser optodes allowed for data collection at four wavelengths (785, 808, 830, and 850 nm) with a sampling frequency of 4.8 Hz. FNIRS data were acquired through NIRStar software (NIRx Medical Technologies LLC, Berlin, Germany). Synchronized triggers were sent to the fNIRS and eye-tracking systems from the MATLAB presentation script using a Cedrus C-pod (Cedrus Corporation, San Pedro, CA, United States) to ensure accurate marking of events.

Each participant’s head circumference was measured to select the appropriate fNIRS cap. The cap was carefully fitted by positioning Cz exactly halfway between the nasion and inion, and halfway between the left and right tragus. Experimenters took great care in ensuring appropriate placement of the cap prior to data collection. A loose black plastic cap (similar to a shower cap) was placed over top of the fNIRS cap and optodes to minimize any effects of environmental light sources on the signal.

### Behavioural measures

#### Autism Spectrum Quotient

The Autism Spectrum Quotient (AQ) was completed by participants to assess autistic-like traits (Baron-Cohen *et al*. 2001). Participants responded to items (e.g., “New situations make me anxious”) on a 4-point Likert scale of 1 (definitely agree) to 4 (definitely disagree). In the current sample, participants showed variability in AQ scores (mean = 18.1, SD = 5.2, range = 9-28) and none of the participants scored above the conventional cutoff (32) for clinical likelihood of autism spectrum disorder (Baron-Cohen *et al*. 2001).

#### Eye tracking

Eye tracking data were acquired using an EyeLink 1000 system (SR Research, Toronto, Canada). Using pupil and corneal reflection, monocular eye tracking data were collected at 1000 Hz with an accuracy of 0.15°. At the beginning of each run, gaze was calibrated using a standard nine-point calibration on the stimulus computer. During this calibration, participants fixated on the center of the circle as it moved around the screen. This process was repeated, if necessary, until calibration quality was deemed to be acceptable based on the EyeLink recording software.

### Apparatus and set up

#### Participant set up

Participants sat at a table facing a computer screen with the eye-tracking camera, webcam (Logitech C920), and speaker setup such that they did not obstruct the participant’s view of the screen (Figure 1a). A forehead rest was used to stabilize the head during the task to minimize motion artifacts.

In a separate room, the confederate sat at a table facing a computer screen with a webcam (Logitech C920) and an iPad Pro (Apple, iOS version 14.6) for video streaming. Audio for both participant and confederate used the webcam microphones.

#### Unity application

An in-house application was developed in Unity (Unity Technologies; https://unity.com/) to allow for simultaneous streaming of audio and visual input. This allowed for the recording of all trials and sequential presentation of live and pre-recorded video trials. The application also presented the scripts with the stories and question prompts to the confederate. Event timings, presentation of stimuli, and calibration and triggering of the fNIRS and eye tracking systems were all controlled using an in-house MATLAB script calling the Psychophysics Toolbox (Kleiner, Brainard, and Pelli 2007).

### Pre-processing and statistical analyses

#### fNIRS pre-processing

Pre-processing was performed using the Brain AnalyzIR toolbox (Santosa *et al*. 2018) in MATLAB (R2022b; The MathWorks Inc., Natick, Massachusetts, USA), with custom in-house scripts. Data were modelled using an auto-regressive iteratively reweighted least squares (AR-IRLS). With the field of fNIRS growing rapidly, there is little consensus on a standardized pipeline (Yücel *et al*. 2021). Nevertheless, there are multiple advantages to using AR-IRLS instead of the commonly employed ordinary least squares (OLS) models.

Specifically, the AR-IRLS model employs an auto-regressive filter to simultaneously account for serially correlated errors and uses robust weighted regression to iteratively down-weight outliers in the data (Barker, Aarabi, and Huppert 2013). This method has been shown to minimize type I errors and improve performance of the GLM overall, especially when there are motion artifacts in the data (Barker, Aarabi, and Huppert 2013). Because AR-IRLS downweights noisy data points (such as movement-related spikes), it minimizes the need for user-specified selections for the various steps in more traditional pre-processing pipelines, such as arbitrary parameters for motion correction. Based on best practices, as well as our own comparisons across multiple fNIRS data sets, AR-IRLS with minimal pre-processing was more robust than the OLS with varied steps for pre-processing.

Raw data were converted to optical density and then to concentrations of oxygenated (HbO) and deoxygenated (HbR) hemoglobin using the Modified Beer-Lambert Law with a pathlength factor of 0.1. Data were down sampled to 1 Hz for most of the analyses, though for deconvolution analyses down sampling was performed after the group-level GLM because, for this question, the timing of evoked responses was of particular interest. Downsampling reduced the number of time points to one per second, substantially decreasing computational demands while preserving the pattern and robustness of the results observed with higher-resolution data. Data from short-distance channels (SDCs) were included as regressors in the models (all SDCs were regressed from each of the long-distance channels), and this was done independently for HbO and HbR signals.

#### fNIRS task analysis

Data were analyzed in three ways, all utilizing AR-IRLS with short-channel regression.

First a region-of-interest (ROI) approach was used to assess fNIRS activation in three independently defined ROIs: left TPJ (lTPJ), right TPJ (rTPJ) and mPFC. ROIs were defined by data collected from the two ToM localizers and from a Neurosynth (https://neurosynth.org/) meta-analysis for the term “theory mind” (Yarkoni *et al*. 2011). The Neurosynth map (association test at default threshold) was registered to the Colin27 MNI template and channels of the montage were weighted based on likelihood of covering the ROIs. Further details are provided in Results (See *Localizer- and Neurosynth-defined regions of interest)*.

Taking advantage of the event-related task design, for each ROI, deconvolution analysis (using finite impulse response, FIR, functions) allowed for estimation of the time course of activation during the multiple trial phases (cue, story, question, answer). Because the hemodynamic response function (HRF) is less well characterized with fNIRS (as compared to fMRI), the deconvolution method was beneficial as it meant there was no assumption about the shape of the HRF. Brain AnalyzIR toolbox functions were adapted in-house to reduce the computational resources required to calculate the robust random effects modelling on entire deconvolution datasets at native resolution (Santosa *et al*. 2018).

Secondly, we performed an *a priori* contrast (two-tailed paired-samples *t* test) to test whether we could replicate the results from previous fMRI studies that found higher activation for pre-recorded than live humans, but which did not include an avatar condition (Rice and Redcay 2016, Rice, Moraczewski, and Redcay 2016). Following the deconvolution pipeline as described above, the average mean, variance, and covariance were calculated for both human conditions at the peak response of the story period (time points 5-7).

Finally, a channel-wise analysis was used to search for differences elsewhere in the full montage, beyond just the selected ROIs. Activation was modelled by convolving a boxcar function for the full trial duration (16 s) for each of the four conditions. A 2×2 ANOVA was conducted on the resulting beta weights. Full results for channel-wise analyses are included in the supplementary materials.

#### Eye tracking analysis

Video recordings from all trials were saved to allow for gaze data analysis. EyeLink data files were converted to MATLAB files using the Edf2Mat Toolbox (https://github.com/uzh/edf-converter). A custom facial feature detection script using MATLAB dynamically identified four screen zones for the analysis, accounting for any movement of the confederate: eyes, mouth, rest of face, and other parts of the screen. For each condition, time spent looking within each of the screen zones was calculated. Because Memoji faces are distorted, particularly by eyes that are rendered at considerably larger sizes than in a real person, looking times in each zone were normalized by the number of pixels within the zone. A repeated-measures ANOVA (using JASP version 0.18.3) was run for each of the four screen zones and *p* values were considered significant after correcting for the number of tests.

## Results

Across all results, only effects that reached statistical significance (*p* < .05) or showed trends (p < .1) after corrections for multiple comparisons are reported, unless explicitly stated.

### fNIRS localizer results

#### Partly Cloudy localizer

Results from the group-level random effects AR-IRLS GLM for the *Partly Cloudy* localizer on all participants (*n* = 25) indicated increased HbO and decreased HbR for mental versus pain events in three channels over the right TPJ (*p* < .05) (Figure 2a). There were also four significant channels covering the PFC, three corresponding to the expected location of mPFC (*p* < .05). Surprisingly, there were no statistically significant effects for the mental versus pain contrast in channels expected over left TPJ. This contrast was uncorrected for multiple comparisons, as it was only used for the identification of channels for independent ROI analyses.

**Figure 2.**
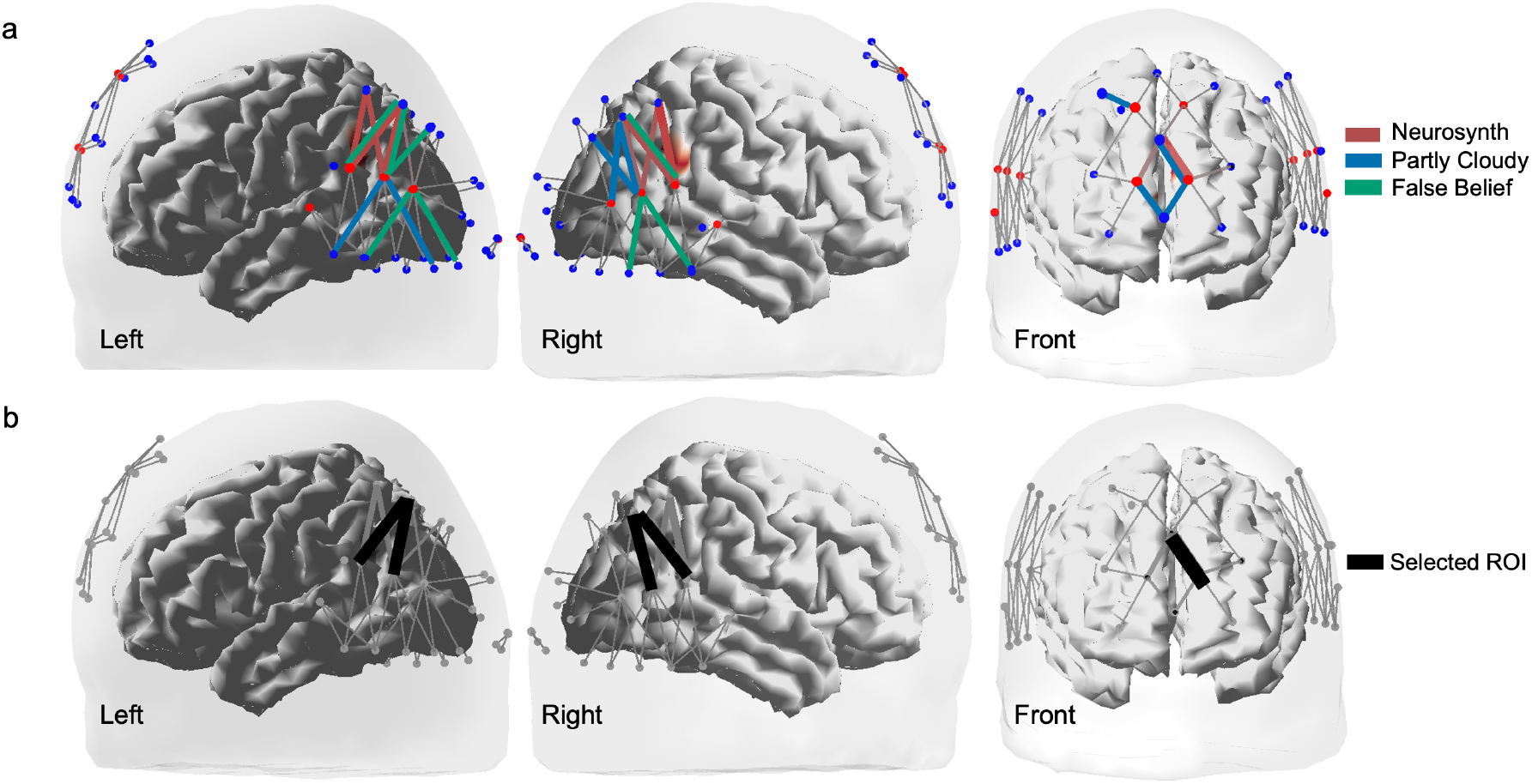
Channel selection using localizer and Neurosynth data. (a) Channels covering left TPJ, right TPJ, and mPFC that showed increased oxygenated hemoglobin (HbO) for mentalizing events (blue, *Partly Cloudy* and green, false belief) or channels that were selected based on Neurosynth meta-analysis for the term “theory mind” (red) were plotted. Using both the localizers and Neurosynth to inform selection, channels were chosen for the region-of-interest analysis if there was convergence from at least two of the methods. (b) This resulted in two channels over left TPJ, two channels over right TPJ, and one channel over mPFC. Homotopic channels for bilateral TPJ were identified as the ROIs, while the mPFC ROI channel was over the left PFC. **Alt text:** Left, right and front views of a head-and-brain rendering showing the fNIRS montage. Top row: channels flagged by Neurosynth (red), the Partly Cloudy localizer (blue) and the false belief localizer (green), clustered over bilateral temporoparietal junction and medial prefrontal cortex. Bottom row: the five channels selected as regions of interest, in black.

#### False belief localizer

A subset (*n* = 16) of the total analytic sample also completed the false belief task. Results from the group-level random effects AR-IRLS GLM found several channels with increased HbO for the false belief versus false photo contrast. Three channels in the right hemisphere and five channels in the left hemisphere covering temporo-parietal regions indicated increased HbO responses (*p* < .05) (Figure 2a). This contrast was also uncorrected for multiple comparisons.

### Region-of-interest results

#### Localizer- and Neurosynth-defined regions of interest

Channels were included in ROI selection if at least two of the three methods converged (i.e., results from either localizer and Neurosynth). This resulted in two channels over left TPJ, two channels over right TPJ, and one channel over mPFC being identified for later ROI analyses (Figure 2b).

#### Deconvolution analysis results

Deconvolution analyses using a robust random-effects AR-IRLS GLM in ROIs were used to general event-related time courses throughout the phases of the trial (Figure 3). A series of repeated-measures ANOVAs, one per 1-s time point, were conducted to evaluate whether the neural responses in left and right TPJ and mPFC were influenced by the realism and/or whether the social interaction was live or pre-recorded. Using FDR correction (Chen *et al*. 2008), we corrected for multiple comparisons for the number of time points and factors in the ANOVA (two main effects and one interaction). Time points that showed significant interaction effects were further evaluated using *post hoc* two-tailed paired-samples *t*-tests. No correction for multiple comparisons was applied for the *post hoc* tests, as the stringent correction of the ANOVAs significantly limited the likelihood of chance interactions (Rosenthal and Rosnow 1991).

**Figure 3.**
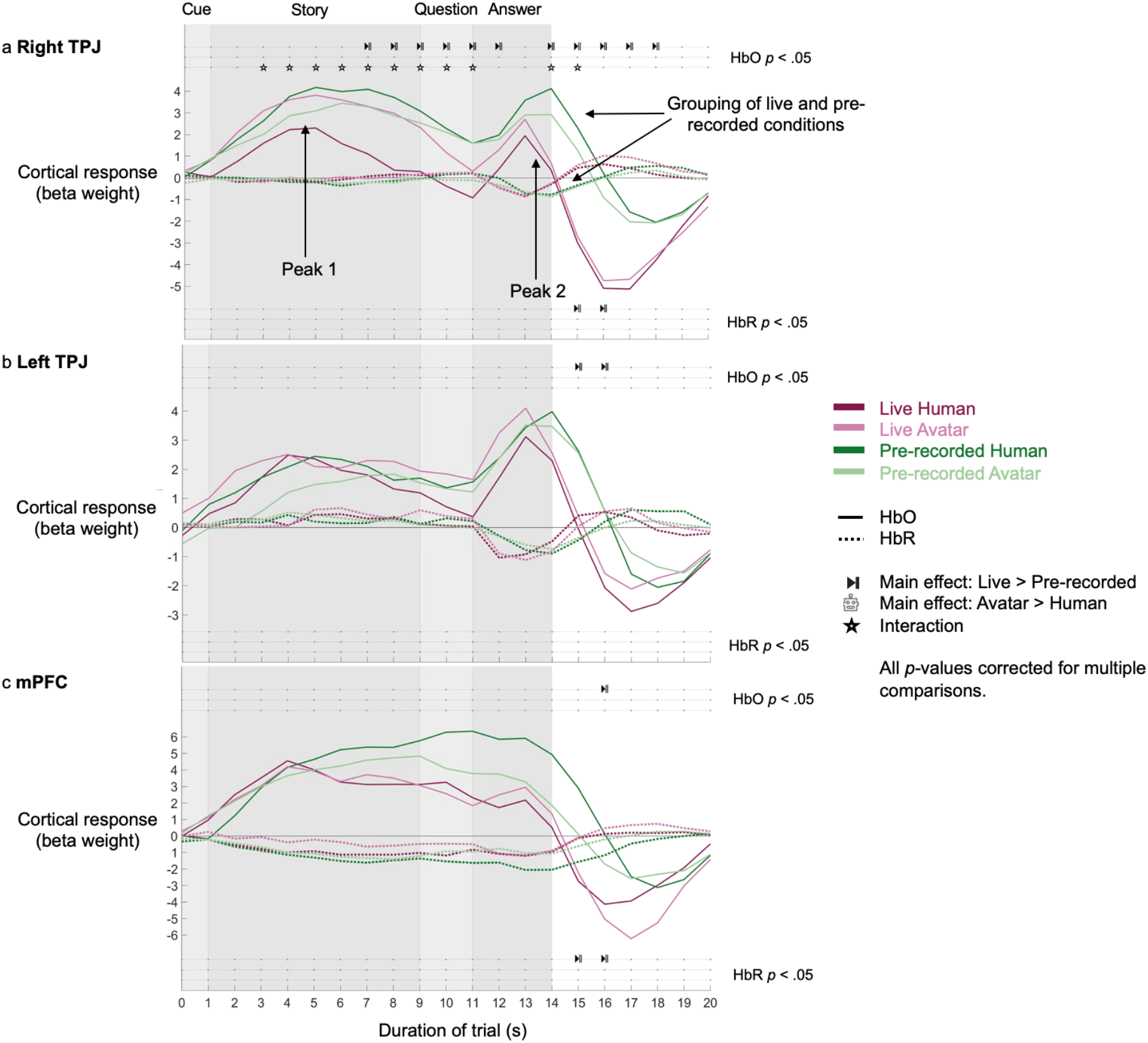
Random effects of deconvolution AR-IRLS GLM. Group-level hemodynamic time courses for three regions of interest (ROIs): (a) right TPJ, (b) left TPJ, and (c) mPFC following random effects AR-IRLS deconvolution. Each of the conditions show distinct peaks corresponding to (1) the story phase and (2) the question phase of the trial, as indicated in panel (a). At each time point of the trial (down sampled to 1-s resolution) an ANOVA was performed to test two main effects and an interaction. Icons represent time points at which the main effects and/or interactions were statistically significant after FDR correction (*p* < .05). Notably, in all three ROIs, after the answer period, the HRF for live conditions declined more quickly than pre-recorded conditions. The interaction for right TPJ during the story phase was explained by a response that was weakest and briefest for the live human condition compared to the other three conditions. Abbreviations: HbO – oxygenated hemoglobin, HbR – deoxygenated hemoglobin. **Alt text:** Three line graphs (right TPJ, left TPJ, mPFC) plotting cortical response across twenty seconds of trial time, with the cue, story, question and answer phases shaded to divide the trial. Each shows changes in oxygenated (solid) and deoxygenated (dotted) haemoglobin across the four conditions (live human, live avatar, pre-recorded human and pre-recorded avatar). Rows of symbols above and below mark time points of significant effects.

#### Deconvolution plot observations

The deconvolution time course plots (Figure 3) reveal several notable features. Deconvolution produced clean time courses with the expected properties: Responses for each condition started and ended with activation levels near zero; the HbO and HbR signals showed the expected anticorrelation; and the timing of responses was consistent with the expected 4-6 s delay in hemodynamic responses. These were important “sanity checks” considering that rapid event-related designs with deconvolution have been uncommon in fNIRS analysis compared to fMRI, where signals may have less noise contamination. Time courses indicated that the HbO signal had distinct responses to different phases of the trial, with two peaks corresponding to the story and question periods. The HbR signal was largely inverted and weaker in magnitude compared to the HbO signal, as expected, with a clearer negative response to the answer phase than the story phase. These results were largely consistent across conditions. In the mPFC, the overall hemodynamic response appeared to be largely sustained for the duration of the trial.

More interestingly, visual inspection of the deconvolved time courses (Figure 3) revealed differences between the four conditions, most notably differences in the duration of activation during the answer phase, with more prolonged activation for pre-recorded than live conditions across all three ROIs. Statistical testing with FDR-corrected ANOVAs at each time point confirmed and clarified these observations (see symbols for each time point in Figure 3). The difference between pre-recorded compared to live conditions (main effect) at the end of the answer period was statistically robust for both HbO and HbR in rTPJ and mPFC and for HbO only in lTPJ. These effects were modulated by an interaction in rTPJ for HbO in which the prolongation for pre-recorded trials was greater when the stimulus was a human compared to an avatar. Significant interaction effects in HbO were also found for the rTPJ throughout the story response, driven by lower activation for the live human compared to the other three conditions. Across all ROIs, there were no significant main effects for visual realism (avatar versus human) in HbO or HbR.

#### Planned comparison of live versus pre-recorded human conditions

Recall that one of our main goals was to test whether current fNIRS results replicated the fMRI effects reported by Rice and Redcay (2016). As such we also performed a planned comparison between the live and pre-recorded human conditions for the peak activation during the story phase (timepoints 5-7 of the deconvolution plots). Analyzing this as a planned comparison avoided the conservatism of the FDR-corrected ANOVA, reducing the risk of Type II errors. Paired-samples *t*-tests indicated that the only significant result for the live versus pre-recorded human contrast was in the HbO response of the rTPJ (*t*(23) = -2.74, *p* = .01). Surprisingly, in our fNIRS data, the live human condition had a *lower* response compared to pre-recorded conditions, a result that goes in the opposite direction to that expected from the fMRI study, which found higher activation during the story phase for live versus pre-recorded. Tests in all other regions and chromophores were non-significant (*p* > .05).

### Channelwise analyses

Channelwise ANOVAs contrasting HbO activation throughout the full trial period are shown in Supplementary Materials and Supplementary Figure S2. Results were largely consistent with ROI analyses, particularly finding stronger activation for pre-recorded versus live conditions and to a lesser degree, for human versus avatar conditions. Effects were more widespread, particularly for channels posterior and ventral to TPJ, and did not align perfectly with channels used to define ROIs. Interestingly, some of these channels may have overlapped with the expected location of face-selective areas such as the posterior superior temporal sulcus and occipital face area.

### Eye tracking results

As shown in Figure 4, participants spent more time looking at the eyes for the human than the avatar, particularly in the live conditions; conversely they spent more time looking at the mouth for the avatar than the human. Specifically, an ANOVA on time spent looking at the eye region showed significant main effect of realism (human versus avatar) for the eye region (*F*(1,19) = 7.96, *p* = .011, η² = .11), with a trend toward an interaction (*F*(1,19) = 3.30, *p* = .09, η² = .03). An ANOVA on the time spent looking at the mouth region showed a significant main effect of live versus pre-recorded (*F*(1,19) = 20.58, *p* < .001, η² = .23). No significant differences in looking time were found for the rest of the face or other parts of the screen.

**Figure 4.**
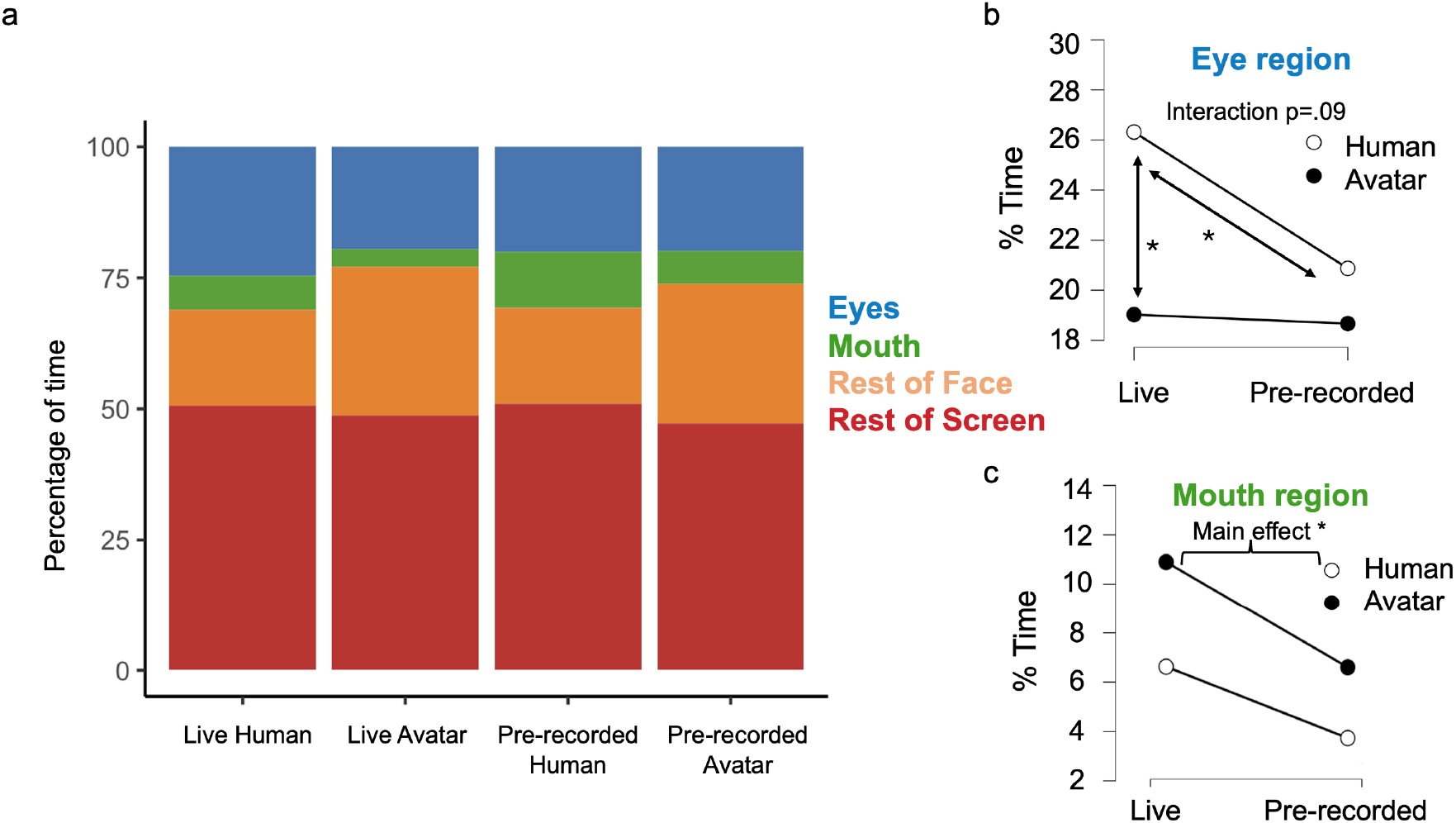
Eye tracking results. (a) Average percentage of time spent looking at each area of interest (eyes, mouth, rest of face and rest of screen) across conditions for the duration of the trial. (b) For the eye region, more time was spent attending to the eyes in the human condition, particularly when the human was live. (c) For the mouth region, more time was spent looking at the mouth during live conditions, especially when it was an avatar. **Alt text:** a) Stacked bar charts of viewing time across eyes, mouth, rest of face and rest of screen for each of the four conditions. (b, c) Line plots of time spent on the eye region and the mouth region, live versus pre-recorded, with separate lines for human and avatar. Line plots emphasize interaction effect for the eye region (b) and a main effect of liveness for the mouth region.

### Autism Quotient results

Correlations between AQ scores and HbO responses to the task were tested for bilateral TPJ and mPFC ROIs. Given the number of tests conducted, none of the results survived correction for multiple comparisons; however, several trends are noteworthy. There was an uncorrected positive relationship between AQ and the HbO response to the live human condition in the lTPJ (r = .40, p = .04), and a similar trend in the rTPJ (r = .36, p = .08), suggesting that individuals with higher autistic-like traits may show greater neural engagement during live human interactions. A trend towards a positive relationship between AQ scores and HbO responses was evident across both TPJ ROIs during the story phase specifically (lTPJ: r = .38, p = .07; rTPJ: r = .36, p = .08). In addition, a trend towards a negative correlation between AQ scores and gaze directed to the face of the live avatar (p = .09) was seen, indicating that individuals with higher AQ scores spent less time looking at the avatar’s face during live conditions.

## Discussion

This study demonstrates that neural responses within the mentalizing network are affected by whether a screen-based interaction is live or pre-recorded and, to a lesser degree, by whether the partner is a human or an avatar. Gaze patterns also differed: participants attended more to the eyes for humans than avatars – particularly during live interactions – and more to the mouth during live than pre-recorded conditions.

The most striking fNIRS effects emerged from the deconvolution analyses, which revealed distinct peaks for the story and answer phases in both TPJ ROIs. In contrast, mPFC responses were more sustained across the trial, consistent with its broader role in social cognition and self-relevant processing (Amodio and Frith 2006, Grossmann 2013). Notably, neural processing across all three ROIs persisted longer during the answer phase of pre-recorded compared to live interactions (especially for HbO). Because the pre-recorded videos were yoked to the live videos, this difference can not be explained by stimulus confounds. A comparable prolongation of lTPJ activation during the answer phase was subsequently observed in a parallel study with 6-to 11-year-old children (Kent 2025), strengthening confidence that the finding is not spurious. The deconvolution analysis also revealed that HbO responses in the right TPJ during the story phase were weakest during live human interactions relative to the other three conditions. Channelwise analyses (as seen in the Supplementary Materials) were largely consistent with deconvolution findings but additionally revealed more widespread effects posterior and ventral to TPJ, though these analyses show activation averaged across the full trial.

Our findings indicate that video-based social interactions are sensitive to naturalness. Live human interactions evoked the lowest and/or briefest neural activation, presumably because this is the most familiar mode of online communication, potentially reflecting reduced inferential demands during genuine social engagement. The findings that gaze behavior and brain activation were similar but not identical between human and avatar conditions suggests that socially interacting with an avatar invokes many of the same mentalizing strategies and systems as a real person, but with notable differences.

### Differences in Activation and Gaze for Live vs. Pre-recorded Conditions

The deconvolution time courses revealed how neural responses evolve over the course of a social interaction and how the timing of such responses is modulated by condition, demonstrating advantages over conventional fNIRS analysis approaches. The typical approach of generating predictors based on convolving boxcar functions with a default HRF would have missed such timing differences, and block averaging would have produced less clear time courses due to overlapping activation across trials (Aarabi, Osharina, and Wallois 2017). Although fNIRS deconvolution can be vulnerable to noise (Santosa *et al*. 2019, Machado *et al*. 2021), our use of AR-IRLS mitigates this by downweighting noisy observations.

Several mechanisms could account for the differential decline in activation at the end of the answer phase. The changes in blood oxygenation measured by fNIRS are sluggish, introducing some uncertainty about the precise underlying neural events. The delayed signal drop could reflect neural processing that is prolonged, begins later, and/or reaches a lower peak (with features for each of these possibilities visible in the time courses in Figure 3). Regardless of the exact mechanism, these results indicate that the aggregate neural activation (from the combined effects of amplitude and/or duration) was lower under more naturalistic conditions.

Eye-tracking results complemented the neural findings. Participants directed gaze more toward the eyes of a human than an avatar, especially under live conditions – consistent with the eyes being a primary cue to a partner’s attention and emotional state, information that is most relevant during real-time interaction. Increased gaze toward the mouth during live interactions may reflect lip-reading or active monitoring of communicative expression cues.

Eye contact plays a central role in naturalistic, in-person social interaction. During fNIRS hyperscanning, real-time eye-to-eye contact evokes TPJ activation and enhanced cross-brain coherence between the TPJs of two individuals, effects that are eliminated when gaze is directed to a pre-recorded video face (Noah *et al*. 2020). Whether such effects occur during live video is an open question, but there are reasons to expect attenuation, since mutual eye-to-eye gaze is impossible over video: a person looking at the camera appears to their partner to be making eye contact, but cannot simultaneously look at the partner’s face on screen (Bohannon *et al*. 2013). Gaze patterns in video-based encounters may also be shaped by the social constraints that govern prolonged gaze toward a real, live person (Laidlaw *et al*. 2011, Risko *et al*. 2012, Freeth, Foulsham, and Kingstone 2013). The gaze differences we observed may therefore partially account for reduced TPJ engagement during live human interactions, suggesting participants adapt their social understanding strategies to the availability of naturalistic cues. Consistent with this, one study found stronger fNIRS activation in TPJ and mPFC during eye contact with a live human compared to a robot (Kelley *et al*. 2021).

### Differences Compared to Prior fMRI Studies

Contrary to predictions and earlier fMRI work reporting higher activation in the mentalizing network for live than pre-recorded interactions (Rice and Redcay 2016, Rice, Moraczewski, and Redcay 2016), our fNIRS results showed stronger rTPJ responses for pre-recorded than live stimuli. Several factors may account for this discrepancy. First, the previous fMRI studies using a similar (Rice and Redcay 2016, Rice, Moraczewski, and Redcay 2016) employed only audio rather than video interactions, though another fMRI study from the same group examined other tasks over video and also found higher activation for live than pre-recorded scenarios (Redcay *et al*. 2010).

Second, and more critically, those studies used pre-recorded stimuli for both conditions while manipulating participants’ beliefs about liveness, whereas our study used genuinely live interactions for both conditions and yoked stimuli across participants. We additionally reinforced awareness of the liveness manipulation by matching the experimenter’s clothing during the in-person meeting to that worn in the live videos. It is therefore possible that subtle differences in participants’ beliefs about the interaction drove the divergent results, consistent with proposals that mentalizing is flexibly engaged as a function of perceived social demands (Schneider, Nott, and Dux 2014) – though the earlier yoked fMRI study using a different task still found higher activation for live conditions (Redcay *et al*. 2010), which argues against this explanation alone. Third, the earlier studies modelled the story phase using standard HRF convolution rather than deconvolution; however, both our deconvolution peak estimates and our standard convolution model of the full trial converged on higher activation for pre-recorded than live conditions, making this an unlikely explanation for the direction of the discrepancy. A fourth possibility is that our study was conducted during the COVID-19 pandemic, which normalised live video interaction, particularly among university student populations; whereas the earlier studies occurred prior to 2020. Familiarity with video communication may have lowered the inferential demands of live video encounters in our sample, consistent with pandemic-related effects on face processing more broadly (Freud *et al*. 2020).

Finally, and most obviously, we used fNIRS whereas the earlier studies used fMRI. Although fMRI and fNIRS results often converge (Cui *et al*. 2011, Pereira *et al*. 2023), direct comparisons of the two techniques have been limited in number and scope, often focusing on motor or sensory paradigms rather than social cognition (for a review see Yang and Wang 2025 Table 3). Thus it is possible that the two techniques could yield different effects. The techniques differ substantially in spatial resolution, temporal resolution, and signal-to-noise ratio. They also differ in the tissue they sample: fNIRS has limited penetration depth and is more sensitive to superficial cortex (gyri) and extracerebral tissue than to deeper structures (sulci), which could contribute to divergent results for regions such as TPJ.

Consistent with previous fMRI findings, exploratory analyses suggested a possible relationship between autistic-like traits and neural responses during live social interaction. Higher Autism Quotient (AQ) scores were associated with increased oxygenated responses in the left TPJ during live human conditions, paralleling reports of AQ-related modulation of activity in the angular gyrus during social interaction tasks (Rice and Redcay 2016).

However, these effects did not survive correction for multiple comparisons and therefore should be interpreted with caution. While not conclusive, these exploratory patterns are consistent with the possibility that left TPJ contributes to individual variability in social-cognitive engagement during real-time interactions.

### Limitations and Future Directions

An important question for future research is what factors of simulated avatars and the social exchange drive differences in mentalizing processes. We deliberately used highly artificial avatars (Memojis) based on the logic of starting with an obvious, pronounced difference. This avoided any potential participant confusion about artificiality and enabled us to show that the attribution of ToM to an agent controlling an avatar was not sufficient to override effects of the visual differences from a real face. However, such synthetic avatars remove many subtle cues to communication that may contribution to the interpretation of social interactions. Most notably, while the avatars show some changes in expression and mouth features with speech, they lack nuances of facial expressions and microexpressions that can communicate emotions and reactions. Notably, even somewhat realistic avatars animated with motion tracking can evoke different fMRI responses to emotion (Kegel *et al*. 2020).

Our avatars were rendered two dimensionally on a flat screen rather than in 3D (Sagehorn 2026) or through virtual reality, which may have an impact. Both the human and avatar in our study did not make eye contact with the participant because of the limitations of screen cameras and the need for the confederate to read their portion of the dialogue aloud.

Technologies are moving toward digital alterations that make indirect gaze at the camera appear as direct gaze toward the partner (e.g., Apple FaceTime Eye Contact); however, at present, many users experience a “creepy” sensation (e.g., Pearson-Jones 2022). As technology is approaching the ability to render ultrarealistic faces that are almost indistinguishable from real faces (Proverbio and Dosaikina 2026) an interesting future question will be the degree to which such realism affects responses. Our results suggest that simply knowing that an avatar is controlled by a real, live human is not sufficient to evoke identical responses to a real human.

Although our choice of paradigm was valuble for comparisons with past fMRI studies (Rice and Redcay 2016, Rice, Moraczewski, and Redcay 2016), the scripted interaction was unnatural. Thus another useful topic for future research would be whether our findings would generalize to more natural and spontaneous tasks. Another interesting direction for future research would be to use hyperscanning to investigate how brain-to-brain synchronization is modulated by the realism of an avatar. One recent study found higher synchronization for visual tracking interactions when the partner was represented by an avatar compared to a cursor (Won *et al*. 2026), though there may also be differences between an avatar and a realistic partner.

### Conclusions

These findings demonstrate that live human interaction elicits gaze patterns and neural signature that are not fully replicated by avatars or pre-recorded video. This has practical implications for the growing use of virtual agents in applied settings, such as education, virtual therapy, and healthcare, where successful interaction depends on mentalizing. Our results suggest that preserving live interactions with real people carries advantages – particularly in domains like education, where there is value in establishing rapport with a teacher and preserving attentional resources for absorbing educational content. While social cognition appears adaptable across real and virtual environments, results suggest that the social brain continues to distinguish between genuine and simulated social partners, even under highly controlled conditions. As technology moves toward increasingly realistic simulations of humans through the generation of compellingly believable avatars under increasing sophisticated control by artificial intelligence, this will no doubt continue to be an important and active area of investigation.

## CRediT author statement

**Michaela Kent:** Conceptualization, Data Curation, Methodology, Software, Investigation, Project Administration, Formal Analysis, Visualization, Writing – Original Draft. **Eva Deligiannis**: Methodology, Investigation, Writing – Review & Editing. **Kevin Stubbs**: Software, Methodology, Formal Analysis, Visualization. **Karsten Babin**: Software. **Emma Duerden**: Conceptualization, Funding Acquisition, Supervision, Writing – Review & Editing**. Jody Culham**: Conceptualization, Methodology, Funding Acquisition, Supervision, Writing – Review & Editing.

## Acknowledgements

We would like to thank Cassia Donga and Lauren MacIntyre for their assistance with data collection, Mozhgan Salimiparsa for assistance with analysis of the eye tracking results, and Homa Vahidi, Joy Hirsch, and Adam Noah for insightful discussions regarding fNIRS data collection and analysis. We also thank the other members of the Culham Lab and Developing Brain Lab for thoughtful feedback throughout the project.

## Data Accessibility

The data that support the findings of this study are available from the corresponding author, JCC, upon request.

## References

Aarabi A, Osharina V, Wallois F. Effect of confounding variables on hemodynamic response function estimation using averaging and deconvolution analysis: An event-related NIRS study. Neuroimage 2017;155(April):25–49. 10.1016/j.neuroimage.2017.04.048.

Aasted CM, Yücel MA, Cooper RJ et al. Anatomical guidance for functional near-infrared spectroscopy: AtlasViewer tutorial. Neurophotonics 2015;2(2):020801. 10.1117/1.nph.2.2.020801.

Adolphs R. The social brain: Neural basis of social knowledge. Annu Rev Psychol 2009;60:693–716. 10.1146/annurev.psych.60.110707.163514.

Amodio DM, Frith CD. Meeting of minds: the medial frontal cortex and social cognition. Nat Rev Neurosci 2006;7(4):268–77. 10.1038/nrn1884.

Bailenson JN. Nonverbal overload: A theoretical argument for the causes of Zoom fatigue. Technol Mind Behav 2021;2(1):1–6. 10.1037/tmb0000030.

Barker JW, Aarabi A, Huppert TJ. Autoregressive model based algorithm for correcting motion and serially correlated errors in fNIRS. Biomed Opt Express 2013;4(8):1366. 10.1364/boe.4.001366.

Baron-Cohen S, Wheelwright S, Skinner R et al. The autism-spectrum quotient (AQ): evidence from Asperger syndrome/high-functioning autism, males and females, scientists and mathematicians. J Autism Dev Disord 2001;31(1):5–17. 10.1023/a:1005653411471.

Bohannon LS, Herbert AM, Pelz JB et al. Eye contact and video-mediated communication: A review. Displays 2013;34(2):177–85. 10.1016/j.displa.2012.10.009.

Cui X, Bray S, Bryant DM et al. A quantitative comparison of NIRS and fMRI across multiple cognitive tasks. Neuroimage 2011;54(4):2808–21. 10.1016/j.neuroimage.2010.10.069.

Dodell-Feder D, Koster-Hale J, Bedny M et al. FMRI item analysis in a theory of mind task. Neuroimage 2011;55(2):705–12. 10.1016/j.neuroimage.2010.12.040.

Freeth M, Foulsham T, Kingstone A. What Affects Social Attention? Social Presence, Eye Contact and Autistic Traits. PLoS One 2013;8(1). 10.1371/journal.pone.0053286.

Freud E, Stajduhar A, Rosenbaum RS et al. The COVID-19 pandemic masks the way people perceive faces. Sci Rep 2020;10(1):22344. 10.1038/s41598-020-78986-9.

Frith CD, Wolpert DM. Decoding, imitating and influencing the actions of others: The mechanisms for social interaction. Introduction. Philos Trans R Soc Lond B Biol Sci 2003;358(1431):431–4. 10.1098/rstb.2002.1260.

Frith U, Frith CD. Development and neurophysiology of mentalizing. Philos Trans R Soc Lond B Biol Sci 2003;358(1431):459–73. 10.1098/rstb.2002.1218.

Glover GH. Deconvolution of impulse response in event-related BOLD fMRI. Neuroimage 1999;9(4):416–29. 10.1006/nimg.1998.0419.

Gobbini MI, Gentili C, Ricciardi E et al. Distinct neural systems involved in agency and animacy detection. J Cogn Neurosci 2011;23(8):1911–20. 10.1162/jocn.2010.21574.

Grossmann T. The role of medial prefrontal cortex in early social cognition. Front Hum Neurosci 2013;7:340. 10.3389/fnhum.2013.00340.

Hamilton AF de C, Brindley R, Frith U. Visual perspective taking impairment in children with autistic spectrum disorder. Cognition 2009;113(1):37–44. 10.1016/j.cognition.2009.07.007.

Hirsch J, Noah JA, Zhang X et al. A cross-brain neural mechanism for human-to-human verbal communication. Soc Cogn Affect Neurosci 2018;13(9):907–20. 10.1093/scan/nsy070.

Hortensius R, Cross ES. From automata to animate beings: The scope and limits of attributing socialness to artificial agents. Ann N Y Acad Sci 2018;1426(1):93–110. 10.1111/nyas.13727.

Jacoby N, Bruneau E, Koster-Hale J et al. Localizing Pain Matrix and Theory of Mind networks with both verbal and non-verbal stimuli. Neuroimage 2016;126(12):39–48. 10.1016/j.neuroimage.2015.11.025.

Kegel LC, Brugger P, Frühholz S et al. Dynamic human and avatar facial expressions elicit differential brain responses. Soc Cogn Affect Neurosci 2020;15(3):303–17. 10.1093/scan/nsaa039.

Kelley MS, Noah JA, Zhang X et al. Comparison of human social brain activity during eye-contact with another human and a humanoid robot. Front Robot AI 2021;7:599581. 10.3389/frobt.2020.599581.

Kent M. From faces to minds: Exploring socio-cognitive development with fNIRS. Western University Open Repository, 2025. https://hdl.handle.net/20.500.14721/31082 (2 Aug. 2026, date last accessed).

Kleiner M, Brainard D, Pelli D. What’s new in Psychtoolbox-3? Perception 36(ECVP Abstract Supplement) 2007.

Laidlaw KEW, Foulsham T, Kuhn G et al. Potential social interactions are important to social attention. Proc Natl Acad Sci U S A 2011;108(14):5548–53. 10.1073/pnas.1017022108.

Machado A, Cai Z, Vincent T et al. Deconvolution of hemodynamic responses along the cortical surface using personalized functional near infrared spectroscopy. Sci Rep 2021;11(1):1–19. 10.1038/s41598-021-85386-0.

Noah JA, Zhang X, Dravida S et al. Real-Time Eye-to-Eye Contact Is Associated With Cross-Brain Neural Coupling in Angular Gyrus. Front Hum Neurosci 2020;14(February):1–10. 10.3389/fnhum.2020.00019.

Okita SY, Bailenson J, Schwartz DL. The mere belief of social interaction improves learning. In: McNamara DS, Trafton JG (eds.), Proceedings of the 29th Annual Meeting of the Cognitive Science Society. 2007, 1355–60.

Pearson-Jones B. The creepy ‘fake eye contact’ feature on Facetime you NEVER knew about: ‘I’m terrified’. Daily Mail. 10 Jul. 2022. https://www.dailymail.com/lifestyle/article-11000627/Apple-iPhone-users-discover-creepy-fake-eye-contact-feature-Facetime.html (2 Aug. 2026, date last accessed).

Pedreño C, Pousa E, Navarro JB et al. Exploring the components of advanced theory of mind in autism spectrum disorder. J Autism Dev Disord 2017;47(8):2401–9. 10.1007/s10803-017-3156-7.

Pereira J, Direito B, Lührs M et al. Multimodal assessment of the spatial correspondence between fNIRS and fMRI hemodynamic responses in motor tasks. Sci Rep 2023;13(1). 10.1038/s41598-023-29123-9.

Petersen SE, Dubis JW. The mixed block/event-related design. Neuroimage 2012;62(2):1177–84. 10.1016/j.neuroimage.2011.09.084.

Pfeiffer UJ, Schilbach L, Timmermans B et al. Why we interact: On the functional role of the striatum in the subjective experience of social interaction. Neuroimage 2014;101:124–37. 10.1016/j.neuroimage.2014.06.061.

Premack D, Woodruff G. Does the chimpanzee have a theory of mind? Behav Brain Sci 1978;1(4):515–26. 10.1017/S0140525X00076512.

Proverbio AM, Dosaikina M. Neural signatures of hyper-realistic AI-generated faces: Dissociating behavioral indistinguishability from implicit neural evaluation. Sci Rep published online 2026. 10.1038/s41598-026-59487-7.

Redcay E, Dodell-Feder D, Pearrow MJ et al. Live face-to-face interaction during fMRI: A new tool for social cognitive neuroscience. Neuroimage 2010;50(4):1639–47. 10.1016/j.neuroimage.2010.01.052.

Redcay E, Schilbach L. Using second-person neuroscience to elucidate the mechanisms of social interaction. Nat Rev Neurosci 2019;20(8):495–505. 10.1038/s41583-019-0179-4.

Rice K, Moraczewski D, Redcay E. Perceived live interaction modulates the developing social brain. Soc Cogn Affect Neurosci 2016;11(9):1354–62. 10.1093/scan/nsw060.

Rice K, Redcay E. Interaction matters: A perceived social partner alters the neural processing of human speech. Neuroimage 2016;129:480–8. 10.1016/j.neuroimage.2015.11.041.

Richardson H, Lisandrelli G, Riobueno-Naylor A et al. Development of the social brain from age three to twelve years. Nat Commun 2018;9(1):1–12. 10.1038/s41467-018-03399-2.

Risko EF, Laidlaw K, Freeth M et al. Social attention with real versus reel stimuli: toward an empirical approach to concerns about ecological validity. Front Hum Neurosci 2012;6:1–11. 10.3389/fnhum.2012.00143.

Rosenthal R, Rosnow RL. Essentials of Behavioral Research: Methods and Data Analysis. 2nd ed., n.p.: McGraw-Hill, 1991.

Sagehorn M. Realism and neural dynamics in face processing: Evaluating electrophysiological markers of cognitive mechanisms across immersive virtual reality and conventional laboratory conditions. Universität Osnabrück, 2026. 10.48693/939.

Santosa H, Fishburn F, Zhai X et al. Investigation of the sensitivity-specificity of canonical- and deconvolution-based linear models in evoked functional near-infrared spectroscopy. Neurophotonics 2019;6(02):1. 10.1117/1.NPh.6.2.025009.

Santosa H, Zhai X, Fishburn F et al. The NIRS Brain AnalyzIR Toolbox. Algorithms 2018;11(5):73. 10.3390/a11050073.

Saxe R, Carey S, Kanwisher N. Understanding Other Minds: Linking Developmental Psychology and Functional Neuroimaging. Annu Rev Psychol 2004;55(1):87–124. 10.1146/annurev.psych.55.090902.142044.

Saxe R, Kanwisher N. People thinking about thinking people: The role of the temporo-parietal junction in ‘theory of mind’. Neuroimage 2003;19(4):1835–42. 10.1016/S1053-8119(03)00230-1.

Schilbach L, Timmermans B, Reddy V et al. Toward a second-person neuroscience. Behav Brain Sci 2013;36(4):393–414. 10.1017/S0140525X12000660.

Schneider D, Nott ZE, Dux PE. Task instructions and implicit theory of mind. Cognition 2014;133(1):43–7. 10.1016/j.cognition.2014.05.016.

Schurz M, Tholen MG, Perner J et al. Specifying the brain anatomy underlying temporo-parietal junction activations for theory of mind: A review using probabilistic atlases from different imaging modalities. Hum Brain Mapp 2017;38(9):4788–805. 10.1002/hbm.23675.

Tomasello M, Carpenter M, Call J et al. Understanding and sharing intentions: The origins of cultural cognition. Behav Brain Sci 2005;28(5):675–91. 10.1017/S0140525X05000129.

Won K, Pillette L, Savalle E et al. Seeing avatars during social interaction in VR may enhance inter-brain synchronization. IEEE Trans Vis Comput Graph 2026;32(7):5641–57. 10.1109/TVCG.2026.3671344.

Yang L, Wang Z. Applications and advances of combined fMRI-fNIRS techniques in brain functional research. Front Neurol 2025;16:1542075. 10.3389/fneur.2025.1542075.

Yarkoni T, Poldrack RA, Nichols TE et al. Large-scale automated synthesis of human functional neuroimaging data. Nat Methods 2011;8(8):665–70. 10.1038/nmeth.1635.

Yücel MA, Lühmann A v., Scholkmann F et al. Best practices for fNIRS publications. Neurophotonics 2021;8(01):1–34. 10.1117/1.nph.8.1.012101.

Yücel MA, Selb J, Aasted CM et al. Mayer waves reduce the accuracy of estimated hemodynamic response functions in functional near-infrared spectroscopy. Biomed Opt Express 2016;7(8):3078. 10.1364/boe.7.003078.

Zhao N, Zhang X, Noah JA et al. Separable processes for live “in-person” and live “zoom-like” faces. Imaging Neurosci 2023;1:1–17. 10.1162/imag_a_00027.

