## Supplemental Figure 1 and 2 for "fNIRS reveals that live social interactions and visual realism influence neural responses"

### Supplementary Materials

#### Montage design

Optode placement for the montage was determined for optimal coverage of the cortical regions of interest. Rather than using a sparse, rectilinear montage, as is common in fNIRS, channels were arranged to provide higher coverage around the regions of interest (Figure S1).

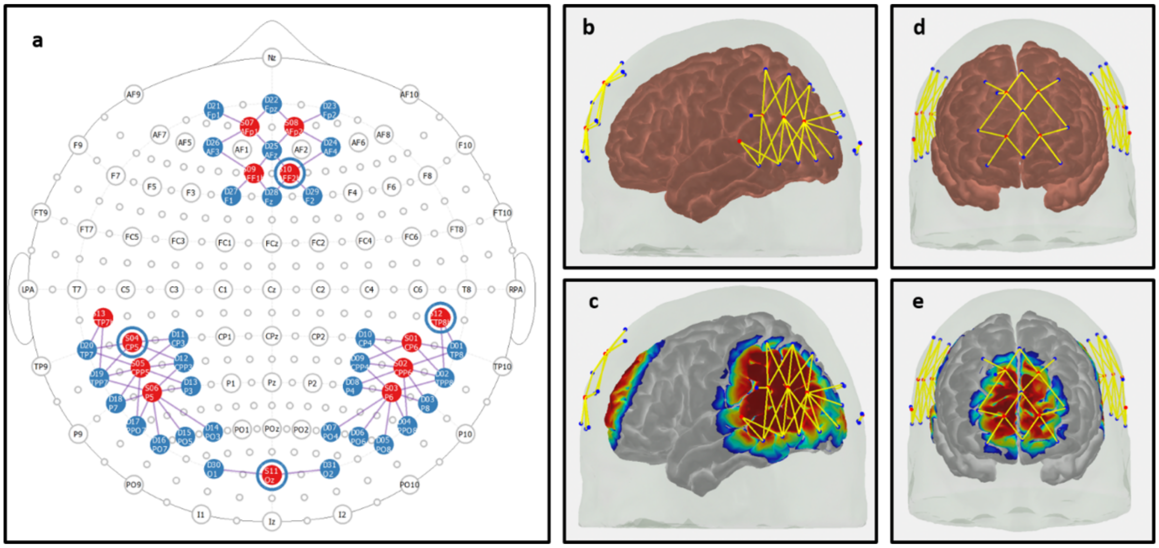


**Figure S1: Custom montage covering the regions of interest.** (a) A 2D representation of the montage layout showing sources (red), detector (blue) and short-distance detectors (blue circles around red sources) locations on the 10-5 system. The custom montage covered mPFC, TPJ, and pSTS (control channels over visual cortex that were not included in the analysis). (b) Left-side view of 3D cortical project of montage, highlighting coverage of left TPJ and pSTS (right and left sides of the montage follow identical layout). (c) Sensitivity modelling performed in AtlasViewer showing montage coverage of left hemisphere. (d) Anterior view of 3D cortical projection of montage targeting bilateral mPFC. (e) Sensitivity modelling of montage coverage over the prefrontal cortex (anterior).

#### Channel-wise results

Channel-wise HbO responses were compared for live and pre-recorded conditions (both human and avatar confederates) across the full trial (Figure S2). Overall, results were similar to those from the ROI-specific analysis, though the effects were more widespread and did not align perfectly with channels expected to cover ROIs.

*Human versus Avatar conditions*

Results from a robust random effects AR-IRLS GLM indicated two channels over left TPJ and two channels over right TPJ that showed increased HbO for the human (live and pre-recorded) versus avatar (live and pre-recorded) contrast (Figure S2). However, only one significant channel per hemisphere overlapped with the localizer- and Neurosynth-identified channels of interest, and the significant channel in the PFC did not overlap with the expected location for mPFC.

*Live versus Pre-recorded conditions*

Results from a robust random effects AR-IRLS GLM indicated widespread decreases in HbO across channels covering the left TPJ, right TPJ, and PFC regions when comparing the live and pre-recorded conditions. For this contrast, many channels appeared to show decreased oxygenated hemoglobin for live compared to pre-recorded conditions, consistent with what was seen in the ROI analysis.

*Interaction effects*

Examining *post hoc* tests for the significant interactions suggested that findings from the channel-wise analysis were largely consistent with the results from the ROI-specific analysis.

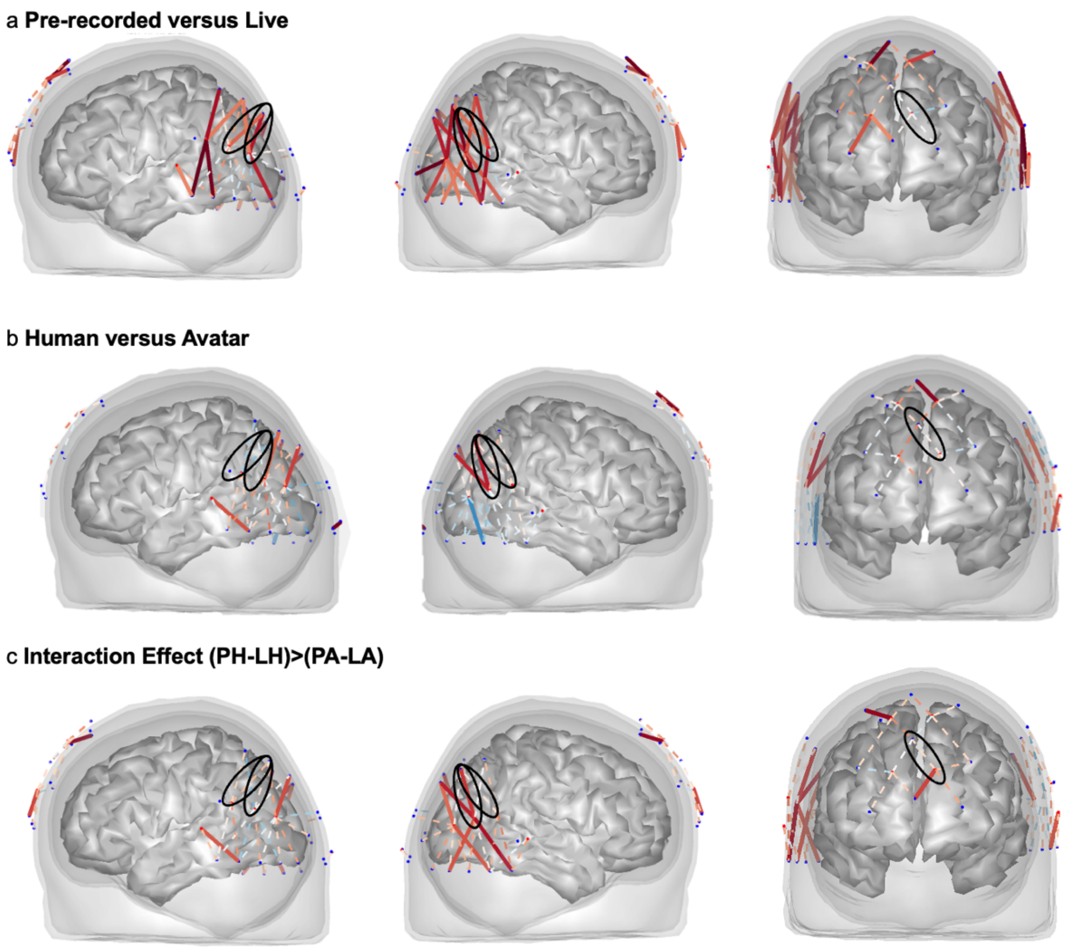


**Figure S2. Random effects of AR-IRLS GLM.** The oxygenated hemoglobin (HbO) results for random effects AR-IRLS across all channels for (a) pre-recorded versus live, (b) human versus avatar, and (c) the interaction effect between the conditions. The convolved GLM was run on the whole trial (cue, story, question, and answer phases). Solid lines indicate channels reaching significance after FDR correction ($q < .05$), with red lines indicating increased oxygenated hemoglobin (HbO) and blue lines indicating decreased HbO for the specified contrasts. Dashed lines indicate channels that did not reach significance. Black circles highlight channels that were used in the ROI analysis. Abbreviations: LH – live human, PH – pre-recorded human, LA – live avatar, PA – pre-recorded avatar, ROI – region of interest.
